# STAPLER: a simple tool for creating, managing and parallelizing common high-throughput sequencing workflows

**DOI:** 10.1101/445056

**Authors:** Jaakko S Tyrmi

## Abstract

STAPLER is a command line program intended for creating, managing and parallelizing bioinformatics workflows. Considerable emphasis has been placed on the ease of adoption and use by effortless installation, simple definition of workflows and quick-start tutorials. Custom workflows can be defined in an easy, modular way allowing the user to choose the desired input data, analysis tools and parameters with a simple parameter file. STAPLER then generates shell scripts that execute the workflow on a personal computer or in a supercomputing environment. Log files are generated to ensure that experimental results can be reproduced, and features are provided for validating run success and allowing rerunning parts of workflow if necessary. STAPLER is freely available on the web at https://github.com/tyrmi/STAPLER, implemented in Python 2 and supported on any UNIX or UNIX-like platform.

## Introduction

Several high throughput sequencing technologies have emerged during the past decade, permitting sequencing on a large scale to answer novel research questions. As it has become easier and faster to obtain huge data sets, handling and processing them has now become one of the most time-consuming task for many investigators. A bioinformatics workflow may include trimming low quality reads and removing adapters or contamination, alignment of reads to a reference genome, editing the aligned reads by removing duplicate reads, variant calling and filtering variants to ensure high data quality for downstream analysis (Pfeifer 2017). Several suitable bioinformatics tools exist for performing each of these steps but chaining the software tools to form a functioning workflow can be tedious.

Workflows can be built by writing simple shell scripts listing necessary command lines for each input file for each step in the pipeline. Such approach is probably quite common among less computationally oriented investigators as it is straightforward and only requires very basic understanding of UNIX scripting. More programming-oriented researchers may also prefer UNIX scripting, as it provides a fine-grained control over the execution of the entire workflow. However, it is also an error prone and tedious approach. When problems occur, it may be difficult to infer which parts of the workflow have failed and how to rerun only the necessary parts. In addition, such hard-coded workflows may be difficult to modify later to incorporate new data and software tools.

Many tools exist to define workflows in more generic way in order to simplify and automate this task. Programs such as bpipe (Sadedin et al. 2012) and Snakemake (Köster and Rahmann 2012) provide users a great flexibility and control over each aspect of pipelines, but programming skills are required to use these tools. Omics Pipe (Fisch et al. 2015) provides an easy way to run built-in best practice bioinformatics pipelines but creating custom pipelines requires Python scripting. Taverna (Oinn et al. 2004) and Galaxy (Goecks et al. 2010) are more beginner friendly as they provide a graphical user interface. They also include powerful features, including the ability to connect to various third-party databases and the possibility of branching workflows. However, it may not always be possible for end users to install such tools to all platforms, especially those provided by a third party, as admin rights or extensive configuration may be necessary. When available, these tools can provide great utility, but many investigators may still want to work in command line environment, particularly if easy-to-use tools are available.

## STAPLER workflow manager

STAPLER (<u>s</u>imple and swif<u>t</u> bioinform<u>a</u>tics <u>p</u>ipe<u>l</u>ine manag<u>er</u>) was developed as an easy to use workflow manager, targeting non-programmer investigators who have basic skillset in working with UNIX command line tools. To provide a quick to learn environment some flexibility of similar workflow tools is absent, such as the ability to branch workflows and interaction with third party databases, yet powerful features are provided to enable many NGS bioinformatics use cases. STAPLER is created as command line toolkit requiring only Python 2.7, which is often included in most UNIX-based operating systems. Therefore, STAPLER installation only requires cloning files from github and editing a configuration file by defining file paths for each bioinformatics software one intends to use. Documentation is provided as a pdf manual included with the software, built in help accessible from the command line and short introductory video tutorials. Example datasets are also included with the software.

Workflows are defined in STAPLER by creating a workflow file including job name, location of input files and a sequence of bioinformatics tools to apply (Listing 1). Also, optional parameters for possible workload manager may be included. A workflow may contain any tool that is supported by STAPLER’s built-in catalogue of bioinformatics software. I addition, STAPLER has the flexibility to allow the use of most tools not found in the catalogue as well. Workflows are modular, so the user may chain tools freely given that the output type of each tool matches the input type of the next tool. The workflow file is then given as an input file to STAPER:

$ python STAPLER.py <workflow_file_path> [options]

STAPLER then generates a shell script file containing all commands necessary to execute the workflow for each input file. Output files of each workflow step are placed into a directory tree providing users an easy access to them. Utilities are also provided to save storage space by compressing intermediate files.

STAPLER enables users to parallelize workflows by in supercomputing environment by providing integration with SLURM, SGE and several other resource managers. Workflows can also be parallelized without resource manager in any environment by running tasks as background tasks. The embarrassingly parallel nature of NGS data sets is utilized in parallelization by executing the workflow of each input file in a separate process. If jobs fail, necessary parts can be rerun from the point of failure.

### 1. Listing

Example input workflow file for STAPLER including raw read alignment to reference using soap aligner (R. Li et al. 2009), alignment processing with picard (http://broadinstitute.github.io/picard/) and samtools (H. Li et al. 2009), and finally variant calling with freebayes (Garrison and Marth 2012) and further filtering with vcftools (Danecek et al. 2011).

|  |
| --- |
| JOB NAME: workflow_1 |
| STARTING POINT DIRECTORY: /path/to/input_directory |
| PROJECT DIRECTORY: /path/to/workflow_1_project |
| COMMANDS: |
| soap --!read_format _R? -D /path/to/reference.fasta |
| Picard_SamFormatConverter -!in_type .sam -!out_type .bam |
| Picard_SortSam -SORT_ORDER coordinate |
| Picard_MarkDuplicates -REMOVE_DUPLICATES true |
| samtools_index |
| freebayes -f /path/to/reference.fasta -u -i -= -X --!input_files_per_command all |
| vcftools --recode --minDP 3 --max-missing 0.7 |
| COMMANDS END: |

Different variations of the same workflow, for instance testing different software tools or their parameters, can be made swiftly by duplicating and modifying the workflow file. Similarly, an existing workflow may easily be applied to another set of input data by minimal edits to the workflow file.

## Summary

Several existing software provide workflow management tools with graphical user interface for ease of use for non-programming biologists. However, installing and configuring these tools may not be possible on all platforms or may require advanced technical skills. Workflow management tools developed for command line offer great control and flexibility over the execution of workflows, but only investigators adept at programming may adopt them. STAPLER is a bioinformatics workflow management tool with emphasis on ease of use, modularity and flexibility for implementing most next generation sequencing workflows. Features are provided for parallelizing jobs in super computer environments, compressing intermediate files and rerunning failed jobs.

## Acknowledgements

I thank members of Plant Genetics Research Group of University of Oulu and especially Tiina Mattila and Komlan Avia for helpful discussions and insightful feedback that helped to improve this software. I also thank Tanja Pyhäjärvi and Outi Savolainen for helpful comments on this manuscript.

